# Wrapping of single-stranded DNA by Replication Protein A and modulation through phosphorylation

**DOI:** 10.1101/2024.03.28.587234

**Authors:** Rahul Chadda, Vikas Kaushik, Iram Munir Ahmad, Jaigeeth Deveryshetty, Alex Holehouse, Snorri Th.d Sigurdsson, Brian Bothner, Reza Dastvan, Sofia Origanti, Edwin Antony

**Affiliations:** Department of Biochemistry and Molecular Biology, Saint Louis University School of Medicine, St. Louis, MO, 63104, USA; Department of Chemistry, Science Institute, University of Iceland, 107 Reykjavik, Iceland; Department of Biochemistry and Molecular Biophysics, Washington University in Saint Louis School of Medicine, St. Louis, MO, 63110, USA; Department of Chemistry and Biochemistry, Montana State University, Bozeman, MT 59717; Department of Biology, Saint Louis University, St. Louis, MO 63103, USA

## Abstract

Single-stranded DNA (ssDNA) intermediates, which emerge during DNA metabolic processes are shielded by Replication Protein A (RPA). RPA binds to ssDNA and acts as a gatekeeper, directing the ssDNA towards downstream DNA metabolic pathways with exceptional specificity. Understanding the mechanistic basis for such RPA-dependent specificity requires a comprehensive understanding of the structural conformation of ssDNA when bound to RPA. Previous studies suggested a stretching of ssDNA by RPA. However, structural investigations uncovered a partial wrapping of ssDNA around RPA. Therefore, to reconcile the models, in this study, we measured the end-to-end distances of free ssDNA and RPA-ssDNA complexes using single-molecule FRET and Double Electron-Electron Resonance (DEER) spectroscopy and found only a small systematic increase in the end-to-end distance of ssDNA upon RPA binding. This change does not align with a linear stretching model but rather supports partial wrapping of ssDNA around the contour of DNA binding domains of RPA. Furthermore, we reveal how phosphorylation at the key Ser-384 site in the RPA70 subunit provides access to the wrapped ssDNA by remodeling the DNA-binding domains. These findings establish a precise structural model for RPA-bound ssDNA, providing valuable insights into how RPA facilitates the remodeling of ssDNA for subsequent downstream processes.

## INTRODUCTION

Replication Protein A (RPA) is an essential eukaryotic single-stranded DNA (ssDNA) binding protein that sequesters transiently exposed ssDNA during various DNA metabolic processes(1-4). RPA binds to ssDNA with high affinity (*K*_d_ ∼10^-10^ M)(5,6), resolves secondary structures, and shields it from nucleolytic degradation(7-10). Notably, the assembly of RPA filaments on ssDNA acts as a trigger for the DNA damage checkpoint(11-14). RPA-ssDNA filaments also function as an interaction hub by recruiting over thirty proteins/enzymes and promoting their assembly on DNA with correct binding polarity(15-17). This multifaceted functionality is enacted through six oligonucleotide/oligosaccharide binding (OB) domains housed within a heterotrimeric complex (**Figure 1A**)(18-20).

**Figure 1.**
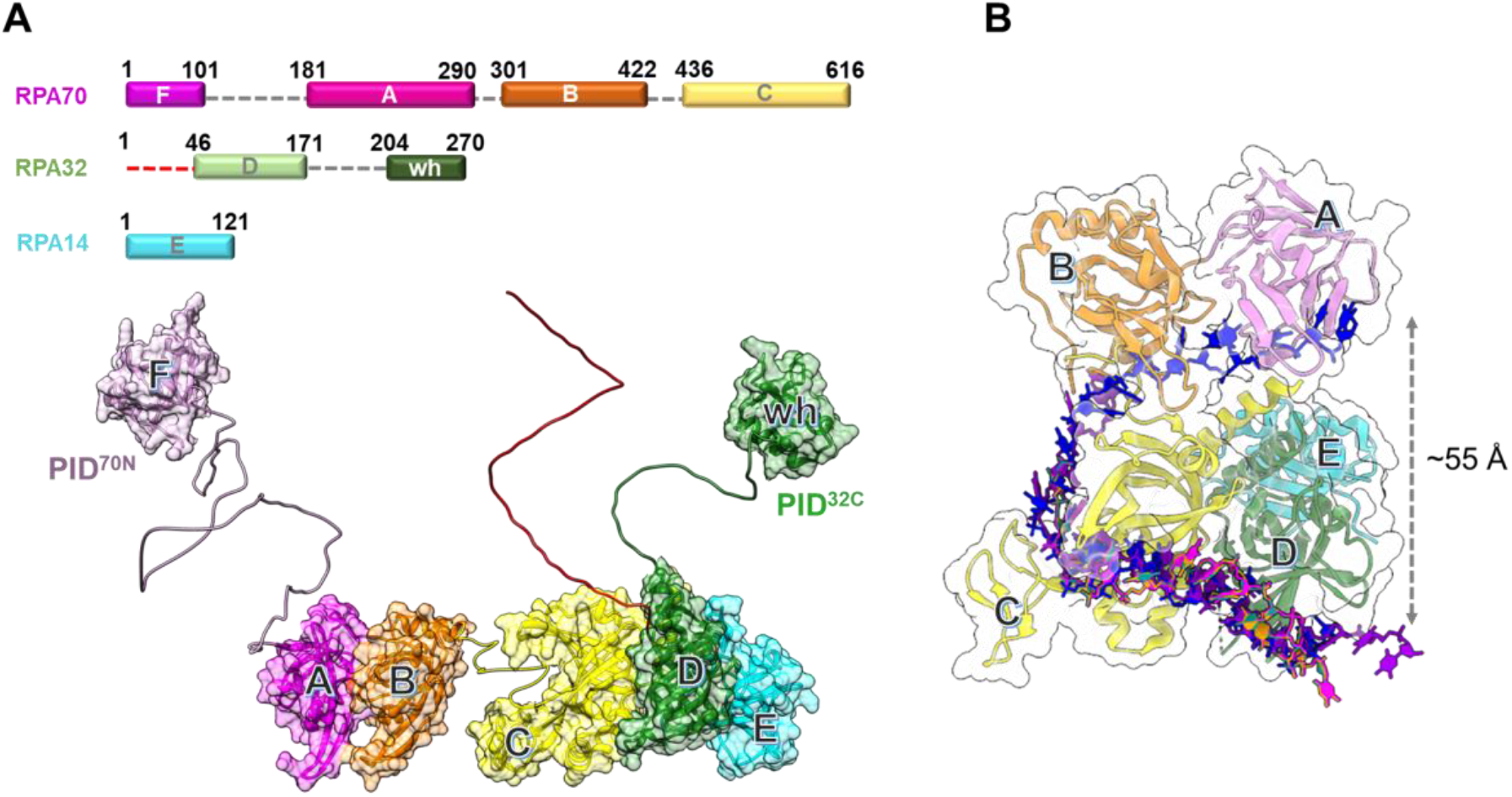
Architecture of the Replication Protein A (RPA) – ssDNA complex. **A)** The three subunits of RPA – RPA70, RPA32, and RPA14 house several oligonucleotide/oligosaccharide binding (OB) domains that are classified as either DNA binding domains (DBDs: DBD-A, DBD-B, DBD-C and DBD-D) or protein-interaction domains (OB-F/PID^70N^ and winged helix (wh)/PID^32C^). A structural model of RPA is shown and was generated using information from known structures of the OB-domains and AlphaFold models of the linkers. **B)** Crystal structure of *Ustilago maydis* RPA bound to ssDNA is shown (PDB 4GNX). This structure lacks OB-F, the F-A linker, wh, and the D-wh linker. ssDNA from cryoEM structures of *Saccharomyces cerevisiae* RPA (PDB 6I52) and *Pyrococcus abyssi* RPA (8OEL) are superimposed and reveal an end-to-end DNA distance of ∼55 Å. Structural data supports partial wrapping of ssDNA around the OB domains.

Human RPA is a constitutive heterotrimer composed of three subunits RPA70, RPA32, and RPA14. Six OB domains (A-F) are spread across the three subunits. OB-F, A, B, and C reside in RPA70 and are connected by disordered linkers. RPA32 harbors OB-D and a winged helix (wh) motif. OB-E is a structural domain in RPA14 and holds the three subunits together as part of a trimerization core (Tri-C) along with OB-C & OB-D(21). Domains A, B, C, & D primarily coordinate ssDNA interactions and are termed DNA-binding domains (DBDs). OB-F and wh coordinate protein-protein interactions and are called protein-interaction domains (PIDs). Dynamic rearrangements of the DBDs and PIDs are observed upon binding to DNA and in response to post-translational modifications(22,23). Each DBD exhibits a moderate affinity for ssDNA binding. Consequently, their collective action engenders a remarkably high affinity and stoichiometric binding of RPA to ssDNA(1). The intrinsic dynamic binding/dissociation characteristics of each DBD transiently unveil pockets within the buried ssDNA, enabling access for incoming proteins while the RPA complex remains attached to the ssDNA(19,24,25).

Two crucial parameters play a pivotal role in unravelling the intricate RPA-ssDNA interactions: a) the length of the ssDNA, and b) the shaping or contour of the RPA-ssDNA complex. The length of the ssDNA determines the number of assembled RPA molecules, subsequently influencing the variety and number of interactors recruited during a specific cellular DNA metabolic process(6). Concurrently, the shape of the complex determines the positioning of the ss-ds junction, or fork, respective to the 3′ or 5′ termini. On average, the occluded site-size for a single human RPA heterotrimer is ∼18-25 nucleotides (nt)(26). Therefore, the density of RPA molecules can be reasonably inferred to be proportional to the length of the ssDNA. Additionally, employing poly-pyrimidine (dT) ssDNA as experimental substrates further mitigates interference arising from secondary structures.

Surprisingly, there is a lack of experimental consensus regarding the configuration of single-stranded DNA (ssDNA) within the RPA-ssDNA complex. In both X-ray and CryoEM structures of diverse RPA-ssDNA complexes, the ssDNA displays a ‘C’ shape, exhibiting end-to-end distances typically ranging between ∼5 to 8 Å (**Figure 1B**). However, the structural findings could be influenced by the utilization of truncated RPA, or a limited subset of its DBDs, alongside potential artifacts originating from crystal packing (crystallography) or dynamics-induced factors (CryoEM). In contrast, biochemical and bulk FRET measurements suggest a complete linearization of even short 30 nucleotide (nt) long ssDNA(19). However, bulk FRET assays are often a readout of multiple underlying populations, especially at relatively high protein to ssDNA ratios.

To mitigate experimental biases or limitations, and accurately determine the contour of ssDNA bound to RPA, we employed solution-based single-molecule confocal FRET and Double Electron-Electron Resonance (DEER) spectroscopy(27-30) to directly measure the end-to-end distance of ssDNA, both in the absence and presence of RPA. On average, we observe only a small <3 nm increase in the ssDNA end-to-end distance in the RPA-bound complex. Our results suggest a partial wrapping of ssDNA around RPA, exhibiting a contour closer to that observed in the crystal structure(18). Therefore, models depicting RPA-ssDNA interactions during DNA metabolism should consider the significant curvature induced in the DNA lattice upon RPA binding. Furthermore, we show that the post-translational modification of RPA through phosphorylation at Ser-384 in the RPA70 subunit introduces substantial changes in the arrangement of the DBDs and PIDs while causing only minimal changes in the pattern of ssDNA wrapping. Thus, access to internal ssDNA segments can be made available by remodeling the RPA domains to serve specific DNA metabolic roles without substantially altering the path of DNA.

## MATERIAL AND METHODS

### Preparation of oligonucleotides

Unlabeled and Cyanine-5 and Cyanine-3 end-labelled poly-(dT) ssDNA oligonucleotides of various lengths (dT)_n_ (n=15, 30, 45, 60, 80, or 97 thymidine bases; Supplemental Table 1) were purchased from Integrated DNA Technologies (Iowa). For the synthesis of the doubly spin-labelled (dT)_22_, and (dT)_50_ oligonucleotides, 2′-aminouridin nucleotides were incorporated at specific sites (Supplemental Table 1) and post-synthetically spin labelled with an isothiocyanate derivative of an isoindoline nitroxide as described(31).

### Purification of RPA and site-specific phosphoserine incorporation

Human RPA was produced using plasmid p11d-hRPA-WT (a kind gift from Dr. Mark Wold, Univ. of Iowa) and purified from *E. coli* as described(32). Briefly, the plasmid was transformed into BL21 (DE3) cells and transformants were selected using ampicillin (100 μg/mL). A single colony was inoculated in 1 L of Luria Broth and incubated at 37°C without shaking for 20-24 hrs. Cells were then grown at 37°C with shaking at 250 rpm, until the OD_600_ reached 0.6 and then induced with 0.4 mM IPTG. Induction was carried out at 37°C for 3 hours. Harvested cells were resuspended in 30 mL cell resuspension buffer (30 mM HEPES, pH 7.8, 300 mM KCl, 0.02% Tween-20, 1.5X protease inhibitor cocktail, 1 mM PMSF, and 10% (v/v) glycerol). Cells were lysed using 0.4 mg/mL lysozyme treatment followed by sonication. The clarified lysate was fractionated over a Ni^2+^-NTA agarose column. RPA was eluted using cell resuspension buffer containing 400 mM imidazole following a 2M NaCl wash to remove and trace non-specifically bound nucleic acid. Fractions containing RPA were pooled and diluted with H_0_ buffer (30 mM HEPES, pH 7.8, 0.02% Tween-20, 1.5X protease inhibitor cocktail, 10% (v/v) glycerol and 0.25 mM EDTA pH 8.0) to match the conductivity of buffer H_100_ (H_0_ + 100 mM KCl), and further fractionated over a fast-flow Heparin column. RPA was eluted using a linear gradient of H_100_–H_1500_ buffers, and fractions containing RPA were pooled and concentrated using an Amicon spin concentrator (30 kDa molecular weight cut-off). The concentrated RPA was fractionated over a S200 column using RPA storage buffer (30 mM HEPES, pH 7.8, 300 mM KCl, 0.25 mM EDTA, 0.01% Tween-20, and 10% (v/v) glycerol). Purified RPA was flash frozen using liquid nitrogen and stored at –80°C. RPA concentration was measured spectroscopically using ε_280_ = 87,410 M^−1^cm^−1^ .

Human RPA-pSer-384, carrying a phospho-serine at position 384 in RPA70, was expressed and purified using genetic code expansion(33,34). Briefly, pRSFDuet1-hRPA-S384TAG and pKW2-EFSep plasmids were co-transformed in BL21(DE3)Δ*serB E. coli* cells and transformants were selected using chloramphenicol (25 μg/mL) and kanamycin (50 μg/mL). A starter culture was prepared by inoculating colonies into ZY-non inducing media (ZY-NIM; Supplementary Table 2A) followed by overnight growth with shaking at 250 rpm at 37°C. 1% of the overnight starter culture was added to ZY-auto induction media (ZY-AIM; Supplementary Table 2B) and grown until the OD_600_ reached 1.5. The temperature was reduced to 20 °C and grown for an additional 20 hours. Cells were then harvested by centrifugation at 2057 xg for 20 mins and cell pellet was resuspended with Pser-cell-resuspension buffer (30 mM HEPES, pH 7.8, 300 mM KCl, 0.02% Tween-20, 1.5X protease inhibitor cocktail, 1 mM PMSF, 10% (v/v) glycerol, 50 mM sodium fluoride, 10 mM sodium pyrophosphate, and 1 mM sodium orthovanadate). RPA pSer-384 was purified as described above for the wild type protein. Typical yields for RPA-pSer-384 are ∼6.8 mg of pure protein/L of culture. pSer incorporation was confirmed using mass spectrometry and analysis on Phos-Tag SDS PAGE.

### Single-molecule FRET measurements

smFRET data were collected on an EI-FLEX bench-top microscope from Exciting Instruments Ltd. (Sheffield, UK) . For measurement, 100 μl of a fluorescent sample droplet was placed onto a no.1 thickness coverslip and excited with alternating 520 and 638 nm lasers at 0.22 mW and 0.15 mW power, respectively. Experiments were carried out in buffer containing 50 mM Tris-acetate, pH 7.5, 50 mM KCl, 5 mM MgCl_2_, and 10% (v/v) glycerol. Lasers were sequentially turned ON for 45 μs for each measurement and separated by a dark period of 5 μs for a total of 40 minutes of acquisition. Fluorescence emission photons from freely diffusing molecules were collected using an Olympus 60X (1.2 N.A.) water-immersion objective, focused onto a 20 mm pinhole. 20 pM of fluorescent sample was suitable for a burst rate of 1 Hz. After passing through the pinhole, the photons were split using a 640 nm long-pass filter, cleaned up using 572 nm and 680 nm band-pass filters, and focused onto respective avalanche photodiodes. The photon arrival times and respective detector were saved in HDF5 data format for offline analysis.

After photoirradiation, the background counts from the buffer were comparable to that of DI water at 2-3 counts per second (cps), whereas typical bursts comprised of 50-100 photons. The photoirradiation of buffer was performed using a 100W LED flood light. The entire set-up (i.e., flood light and buffers taken in covered glass beakers) was placed in a cold-room to minimize heating of samples. Analysis of smFRET data was performed in an Anaconda environment, with Jupyter notebooks, using FRETBursts Python package(35). Single molecule photon emission bursts were identified using a dual channel burst search (DCBS) algorithm as previously described (L =10, and F=45 for both channels). The background estimated from an exponential fit to inter-photon delays greater than 1.5 ms was subtracted. The compensation for spectral crosstalk (a), compensation factor for different detection efficiencies between donor and acceptor channel (g), and compensation for direct excitation (d) for EI-FLEX were estimated as 0.0938, 1.591, and 0.05824, respectively. The same factors for measurements on the Picoquant Micro Time (MT200; Picoquant Inc., Germany) instrument were 0.05, 0.85, and 0.1, respectively.

### Continuous Wave (CW)-EPR and DEER spectroscopy

Continuous Wave (CW)-EPR spectra of spin-labeled ssDNA samples ± RPA were collected at room temperature on a Bruker EMX spectrometer operating at X-band frequency (9.5 GHz) using 2 mW incident power and a modulation amplitude of 1 G. DEER spectroscopy was performed on an Elexsys E580 EPR spectrometer operating at Q-band frequency (33.9 GHz) with the dead-time free four-pulse sequence at 83 K. Pulse lengths were 20 ns (π/2) and 40 ns (π) for the probe pulses and 40 ns for the pump pulse. The frequency separation was 63 MHz. Samples for DEER analysis were cryoprotected with 24% (vol/vol) glycerol and flash-frozen in liquid nitrogen. Primary DEER decays were analyzed using a home-written software (DeerA, Dr. Richard Stein, Vanderbilt University) operating in the Matlab (MathWorks) environment as previously described(36). Briefly, the software carries out analysis of the DEER decays obtained under different conditions for the same spin-labelled pair. The distance distribution is assumed to consist of a sum of Gaussians, the number and population of which are determined based on a statistical criterion.

### Confocal smFRET with alternating-laser excitation (ALEX)

The photons emitted during a transit through confocal volume are called a burst and can be used to estimate FRET efficiency (E).

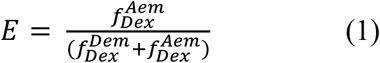

This uncorrected E is a ratio of number of sensitized acceptor emission photons i.e., via energy transfer 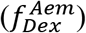 and the sum of number of photons in the donor channel after donor excitation 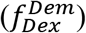 and the number of photons in the acceptor channel after donor excitation 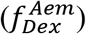. However, to convert the E into distances spectral crosstalk must be taken into account. This especially poses a problem in the low E regimen, i.e. distinguishing low E species from donor alone species is a huge challenge. And this is where alternating laser excitation (ALEX) solves the problem. During an ALEX scheme, both donor and acceptor fluorophores are rapidly excited in alternating fashion. The diffusion coefficient of biomolecules in dilute solutions is of the order of hundreds of μ^2^⁄*s*. And, so typically an alternation frequency of 20KHz is more than sufficient to excite both donor and acceptor fluorophores multiple times during the transit of a single molecule through the confocal volume (∼1 ms). The 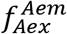 thus estimated can be used to calculate another informative quantity about the population of diffusing single molecules named raw stoichiometry.

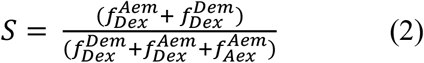

Stoichiometry can be best understood in terms of ratio of total fluorescence photons recorded after donor wavelength excitation to total fluorescence photons recorded after direct donor and acceptor excitation. For donor-only and acceptor-only species the S is close to 1 and 0, respectively. Similarly, the donor and acceptor laser intensity are tuned such that double-labeled molecules scale to an S value of 0.5. Furthermore, binding of protein like RPA to Cy5/Cy3 labelled ssDNA could lead to a reduction in E (i.e. DNA ends are brought further apart), or an increase in S (increase in donor quantum yield due to PIFE) or both. Finally, in order to convert E into distances three addition steps were taken: (1) background intensity in all three photon streams 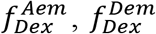, and 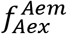 is estimated via mean count rate and subtracted taking the length of burst into account (2) Two, channel cross-talk factors are estimated (a) bleed-through of donor-emission into the acceptor channel, and (b) the direct excitation of acceptor molecules by the donor-laser. These are estimated from the donor-, and acceptor-only molecules in the background-corrected ES histogram, and finally (3) gamma correction, which takes into account the differences in the detection efficiencies of donor and acceptor, their quantum, and the transmission efficiencies of the optical elements etc. The corrected FRET values were converted into distance following the relationship between energy transfer efficiency (E) and donor-to-acceptor separation (r):

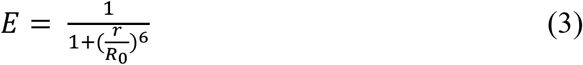

The *R*_0_ or the donor-acceptor distance at which the energy transfer efficiency is 50% was taken to be 5.4 nm. The end-to-end distance for a fully linearized ssDNA molecule was estimated simply by multiplying *l*_*d*_ or rise per base, taken to be 0.67 nm, with the number of the bases in the dT polymer.

## RESULTS

### Single molecule FRET provides an excellent read out of end-to-end ssDNA distances

Single molecule FRET (smFRET) has been used to obtain accurate end-to-end distances for double-stranded nucleic acids (37). Similar measurements of conformational flexibility for ssDNA have been measured using single molecule total internal reflection (smTIRF) microscopy where overhang DNA with varying length of ssDNA were tethered to glass slides (38). Here, we used a benchtop microscope for in-solution smFRET measurements to capture the conformations of ssDNA and RPA-ssDNA complexes (39). First, to test whether this instrumental setup accurately captures the conformational sampling of ssDNA, we measured the changes in FRET as a function of ssDNA length. For these experiments, all ssDNA substrates carried Cy3 (donor) and Cy5 (acceptor) fluorophores at the 3′ versus 5′ ends, respectively (3′-Cy3-(dT)_xx_-Cy5-T-5′; where xx=number of nucleotides). A plot of the FRET efficiency versus length of end-labeled poly(dT) substrates shows an excellent monotonic relationship with high and low FRET captured for (dT)_15_ and (dT)_95_, respectively (**Figure 2**). These measurements obtained using the new benchtop EI-FLEX microscope and methodology are in excellent agreement with similar data collected using a Picoquant MicroTime 200 confocal microscope (**Supplementary Figure 1 & Supplementary Table 3**).

**Figure 2.**
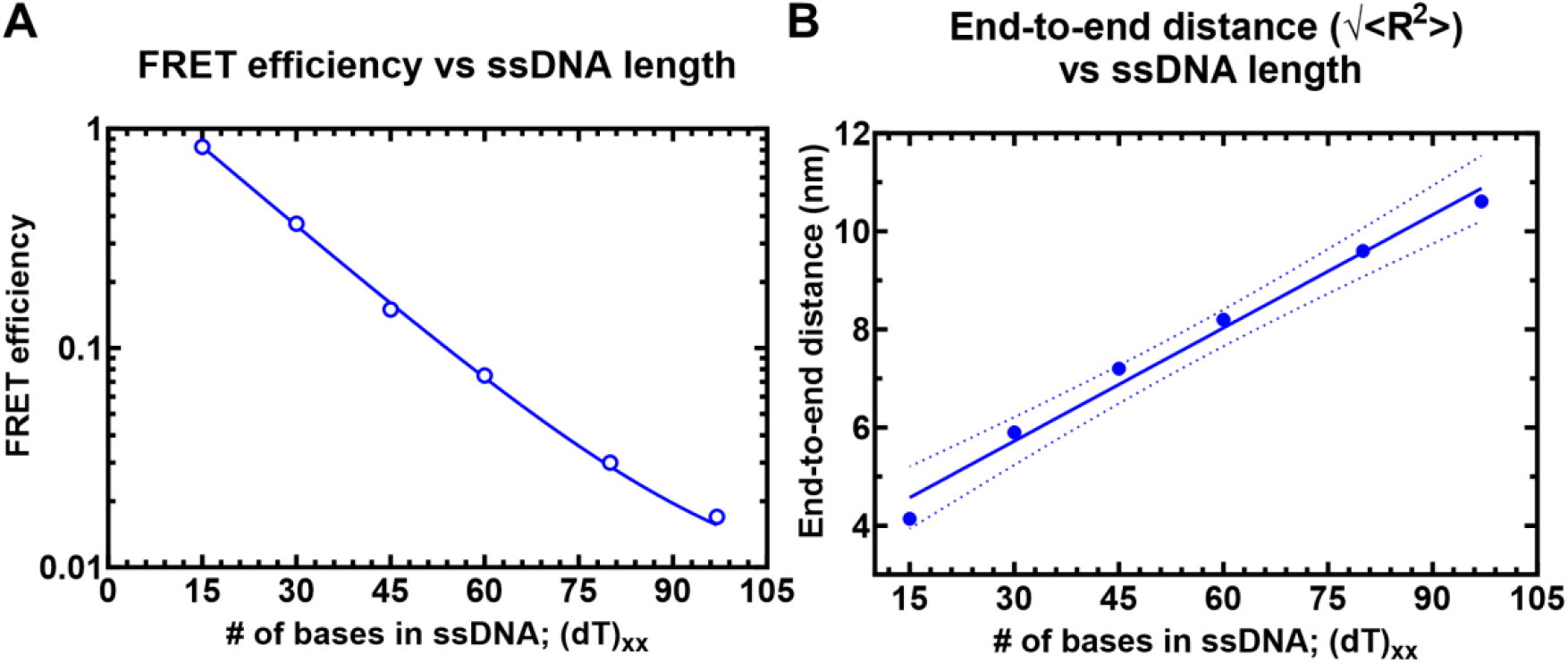
Solution confocal-based FRET measurements accurately report on ssDNA end-to- end distances. **A)** FRET efficiencies were calculated based on single molecule measurements of Cy3 (donor) and Cy5 (acceptor) fluorescence. Data were collected on a series of ssDNA substrates (30 pM) of increasing lengths. Energy transfer events were detected as bursts of photons as the molecules transited the confocal volume. The mean of the distribution is plotted in panel A as a function of length of poly-thymidine (dT_xx_). **B)** The end-to-end distances were calculated based on a R_0_ value of 5.4 nm for the Cy3/Cy5 pair and Eq. 3 described in the Methods.

### RPA binding does not produce a complete linearization of ssDNA

To assess the change in ssDNA shape upon RPA binding, we quantified the FRET changes in the RPA-ssDNA complex. First, in these assays, DNA alone produces a FRET signal proportional to the length (**Figure 2A**). Upon binding to RPA, the FRET signal decreases because of an increase in distance between the donor and acceptor (**Figures 3**). In addition, since the concentration of RPA and ssDNA used in these experiments are low (pM range), we collected data on each ssDNA substrate as a function of increasing RPA concentrations (**Figures 3A-H, Supplemental Figures 2-4, & Supplementary Tables 4-5**). Furthermore, to ensure that we only assessed 1:1 RPA:DNA complexes, we performed parallel mass photometry analysis of the complexes (**Figures 4I-L**). Under all RPA concentrations tested, we captured predominantly stoichiometric RPA-ssDNA complexes. Thus, our interpretation of end-to-end ssDNA distances from the FRET measurements is not influenced by the binding of multiple RPA molecules.

**Figure 3.**
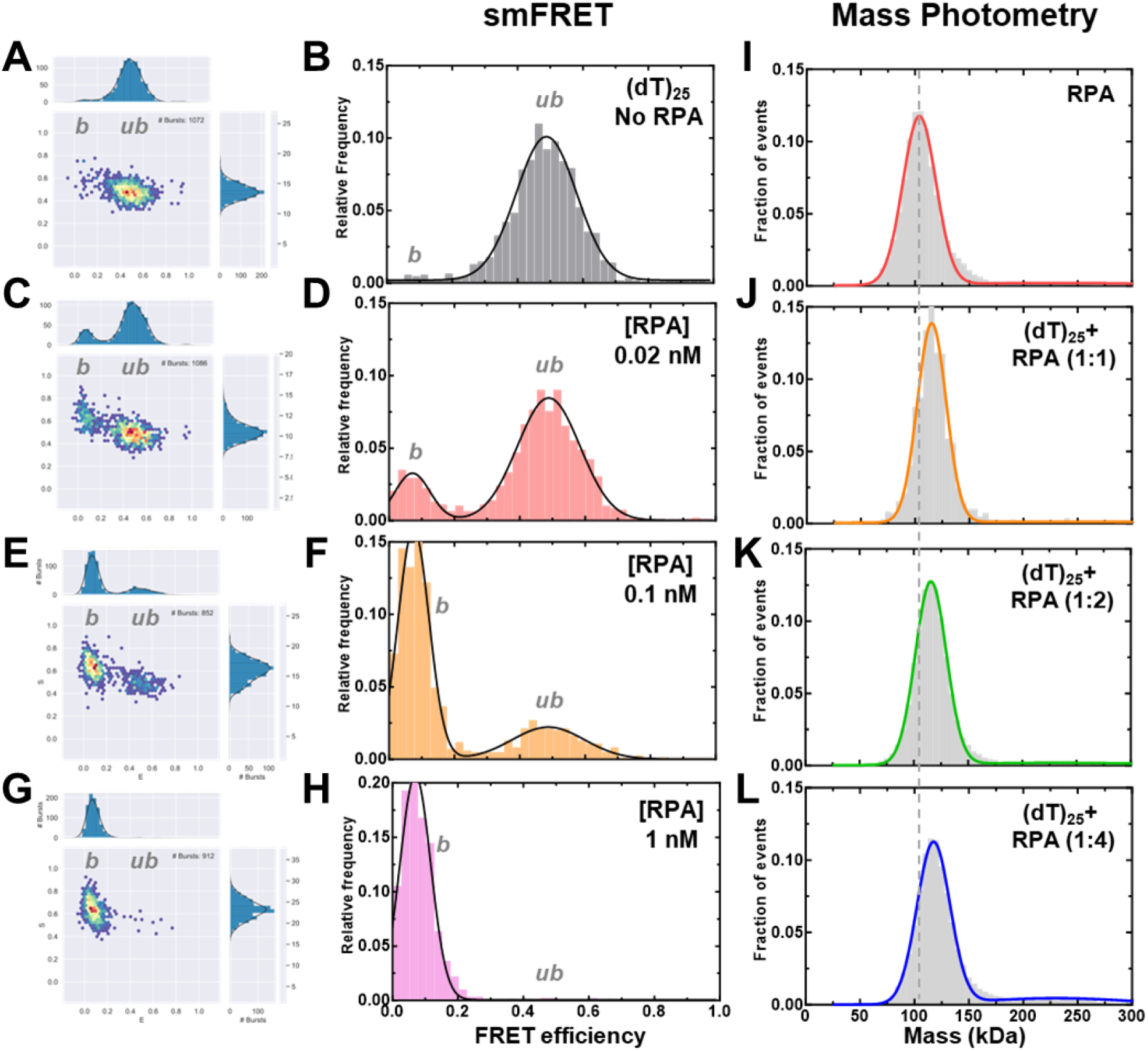
Concentration dependence of RPA-ssDNA complexes. **A-H)** FRET analysis of (dT)_25_ ssDNA bound to increasing concentrations of RPA show a shift from the unbound to bound complex. As RPA concentrations are increased, a complete shift to the bound population is observed. **I-L)** Mass photometry analysis of RPA and RPA-(dT)_25_ complexes show formation of predominantly single RPA bound (dT)_25_ complexes.

**Figure 4.**
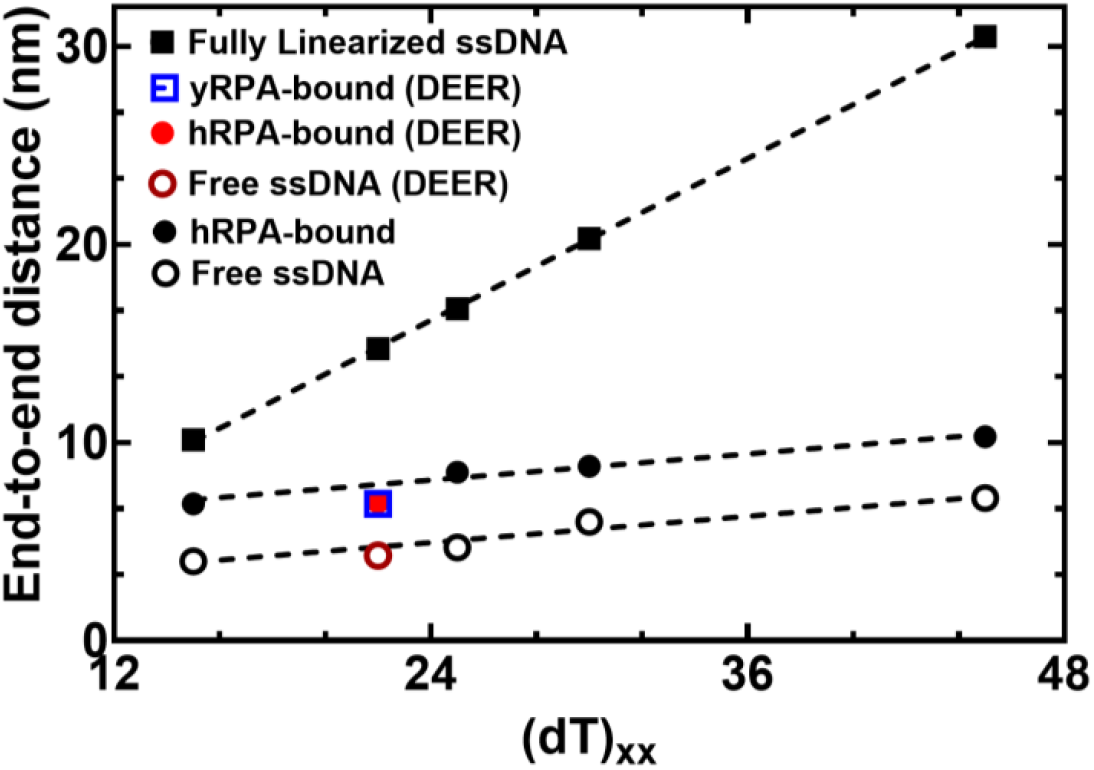
RPA binding produces a modest 2.6 nm increase in end-to-end distance. The estimated distance between 3’ and 5’ end of ssDNA is plotted as a function of nucleotides which make up the chain. Closed squares represent theoretically calculated end-to-end distances assuming the ssDNA was completely linearized. Open and closed circles represent experimental end-to end distance measurements from smFRET analysis ssDNA -/+ RPA, respectively. The figure also shows that end-to-end distances of yRPA-bound, hRPA-bound, and free (dT)_22_ ssDNA, measured using DEER spectroscopy, fall on the respective trend lines suggesting good agreement between experimental measurements performed using two independent biophysical approaches.

smFRET measurements were performed on four ssDNA substrates of varying lengths [(dT)_xx_; xx=15, 25, 30, or 45 nt]. When these measurements are plotted as a function of ssDNA length, the results mirror the data for DNA alone (**Figure 4**). However, for every ssDNA substrate, an overall increase in end-to-end distance of ∼2.6 nm is observed. This surprising result does not agree with the canonical models described for RPA where the DBDs are arranged in a sequential fashion with the ssDNA stretched in a linear manner. Instead, the shorter end-to-end distance measurements better agree with observations in the structural studies where DNA is wrapped around the DBDs, as shown in Figure 1b (18-20). Thus, ssDNA is not linearized, but rather follows the intrinsic curvature of RPA (**Figure 1B**). The second observation is that the 2.6 nm shift in distance is observed across the increasing length of ssDNA. This suggests that DNA occupancy along the curvature of the DBDs is uniform and there is no looping or extrusion of the ssDNA. Finally, the minimal change in distances between the shortest (dT)_15_ and the longest (dT)_45_ ssDNA substrates supports our model that the trimerization core (Tri-C) of RPA contributes most to ssDNA binding stability (and in this case contour or path of wrapping) whereas the F-A-B domains are intrinsically more dynamic.

### DEER spectroscopy confirms wrapping of RPA along the curvature dictated by the DNA binding domains

To obtain another independent experimental validation of our findings from the smFRET analysis, we performed double electron-electron resonance (DEER) spectroscopy. This experiment requires the incorporation of two spin labels, one on each end of the oligonucleotide. 2′-aminouridine was introduced during chemical synthesis of the oligonucleotides and post-synthetically spin-labeled with an isothiocyanate derivative of an isoindoline nitroxide (**Figure 5A & Supplementary Table 1**). The distance between the two spin centers was measured using DEER spectroscopy in the absence and presence of RPA. For a (dT)_22_ substrate, DNA alone produces a broad distribution of distances with two populations and these measurements are excellent agreement with our smFRET data (**Figure 5B** and **Supplemental Table 6**). When RPA is added in equimolar amounts to form a stoichiometric RPA:DNA complex, the end-to-end distance increases to ∼ 7 nm (**Figure 5C**). This measurement is again in very good agreement with our smFRET observations (**Figure 5**). We also checked if this phenomenon was specific to human RPA by performing DEER and smFRET measurements for the *Saccharomyces cerevisiae* RPA-ssDNA complex. On a (dT)_22_ oligonucleotide, the end-to-end distance in DEER measurements was 6.9 nm (**Figure 4**). In smFRET measurements on a (dT)_25_ oligonucleotide, we observed an end-to-end distance of 7.9 nm (**Figure 4**). These distances are similar between the human and yeast RPA, suggesting that the mechanism of ssDNA wrapping is likely conserved across eukaryotic RPA.

**Figure 5.**
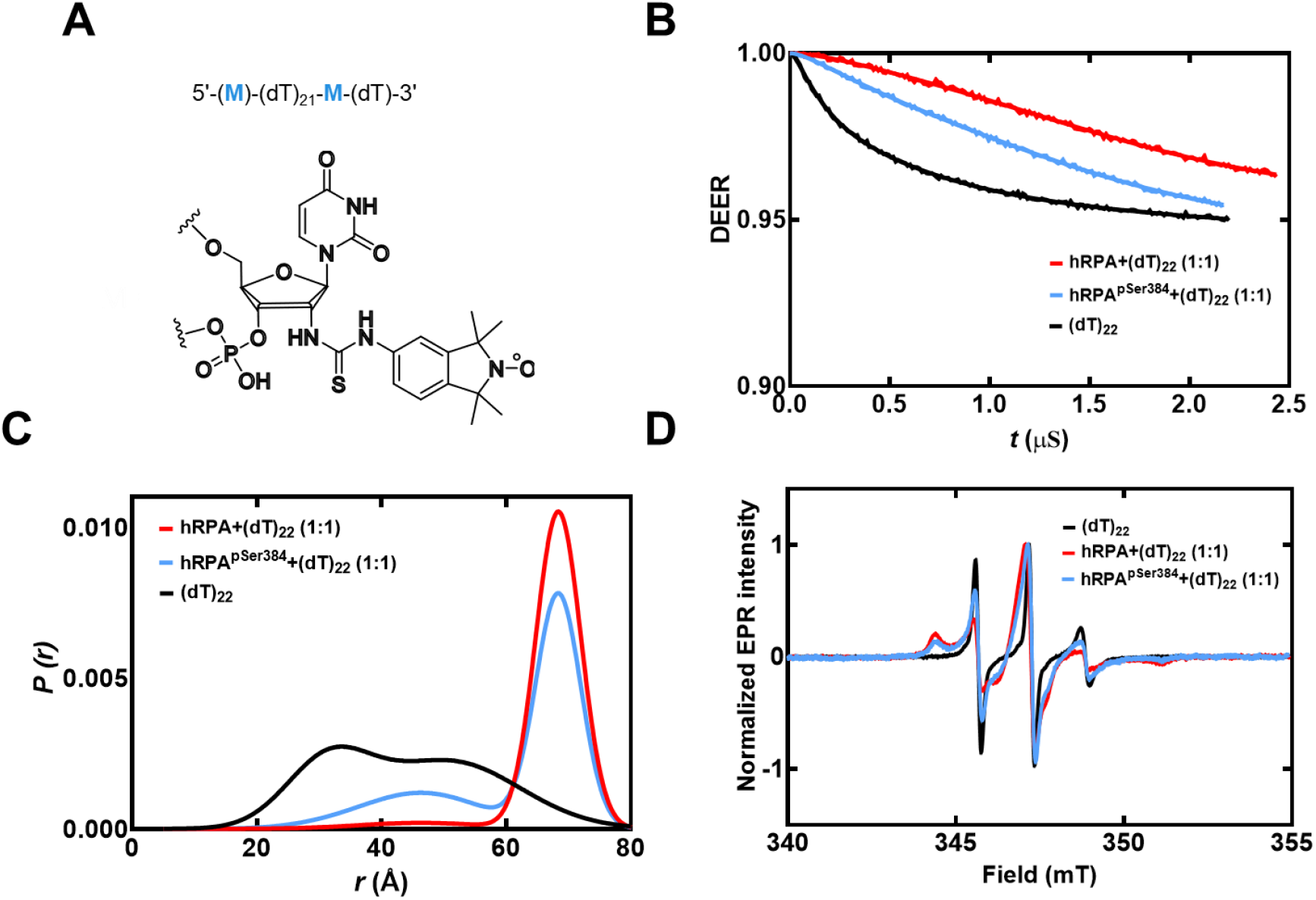
DEER spectroscopy of RPA and RPA-pSer^384^ bound to ssDNA. **A)** Structure of the isoindoline nitroxide spin label and the position on the (dT) oligonucleotide. **B)** Raw DEER decays and fits are presented for the experimentally determined distance distributions *P*(*r*) (**C**). **D)** CW EPR spectra of labeled ssDNA in the absence and presence of RPA and RPA-pSer^384^.

### Computational prediction of the shape of the wrapped ssDNA

In principle, ssDNA should behave as a flexible polymer, in particular when the sequence cannot form secondary structures through intrastrand base-pairing; thus the choice of polyT (38). The dependence of the end-to-end distance for a flexible polymer on the number of monomer units (here nucleotides) generally can be described in terms of polymer scaling laws. We therefore sought to determine if our unbound dT constructs showed characteristic polymer scaling behavior. Fitting the root-mean-squared end-to-end distance against the number of bases reveals extremely good agreement with a polymer scaling law of R_e_ = R_0_N^ν^, where R_0_ is 1.07 nm and ν is 0.50 (**Figure 6A**). This would suggest that under the solution conditions examined, ssDNA (dT) behaves as a flexible chain that conforms to the statistical properties of a Gaussian chain. To further assess the validity of our experimental measurements and conclusions, we compared end-to-end distance obtained previously using a confocal-based smFRET set up by Chen et. al. (40). Gratifyingly, for (dT)_40_ under matching solution conditions (50 mM NaCl), the data from Chen et al. lies directly over the polymer fit generated from our smFRET data (**Figure 6A**).

**Figure 7.**
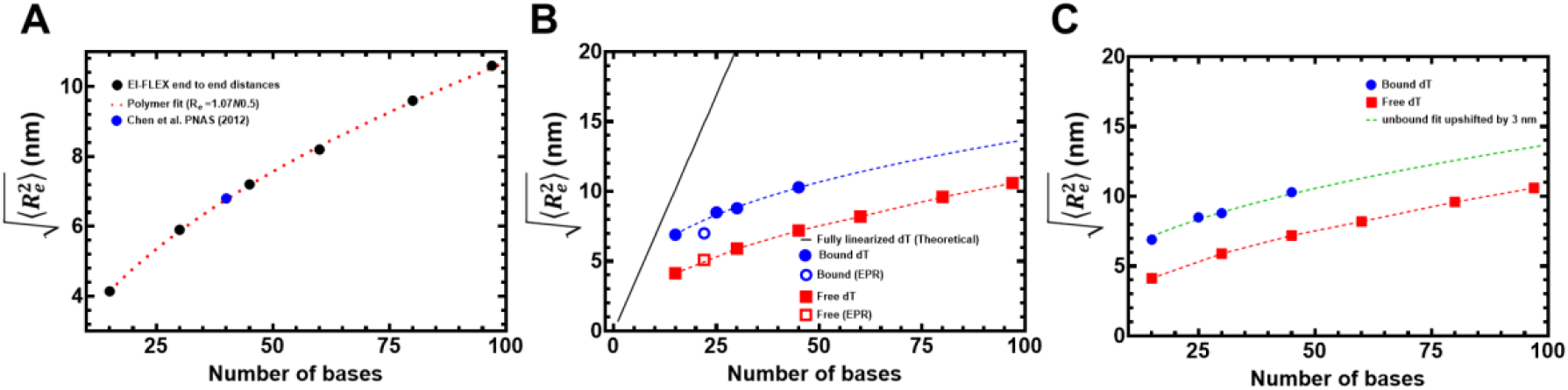
Computational predictions of ssDNA wrapping agree well with experimental measurements. **A)** Black squares are data, and the red dashed line is a fit to those data using a polymer scaling model. The blue circle is data reporting on the end-to-end distance for (dT)_40_ made by Chen et al. 2011). **B)** Blue circles are from RPA-bound (dT) oligonucleotides, red squares are from RPA-free (dT) oligonucleotides. Data include distances measured by both EPR and smFRET experiments. Blue and red dashed lines are polymer model fits to the data, respectively. The black line is the theoretical expected end-to-end distance if the DNA were fully linearized and stretched to its contour length. **C)** The root mean squared end-to-end distance dependence on the number of bases is essentially identical for the bound and unbound states, as highlighted by the fact that the exact polymer fit upshifted by ∼3 nm fully describes the bound dT dependency.

If the bound-state conformation were to form a stretched linear extension, we would expect major deviations from the polymer scaling and an increase in distance between the DNA ends. Instead, bound-state derived distances reveal an identical dependency on the number of nucleotides, shifted by ∼3 nm more expanded than the unbound state (**Figure 6B**). Both bound and unbound distances are far from the trend expected for fully linearized DNA. Taken together, our results suggest the configurational/conformational properties of the RPA-bound ssDNA are relatively similar to unbound, with the exception of a systematic shift in the end-to-end distance, consistent with dT wrapping around RPA in a dynamic state that preserves the DNA’s intrinsic flexibility (**Figure 6C**).

### Post-translational modification of human RPA by Aurora kinase B rearranges the domains without affecting the wrapping of ssDNA

Since ssDNA wraps around the contour of the DNA-binding domains of RPA, we wondered how RPA-interacting proteins might gain access to the buried ssDNA. There are two important features that need to be considered: The first is the intrinsic dynamic interactions between the DBDs and ssDNA, and the second is the configurational rearrangements of the domains with respect to each other and the ssDNA. We, and others, have shown that the domains possess different dynamic properties, with DBD-A and DBD-B being more dynamic on ssDNA compared to DBD-C and DBD-D(19,22-24). Using C-trap experiments, we recently showed that the rates of diffusion for RPA on long stretches of ssDNA are regulated by the trimerization core(41). Thus, incoming proteins can access the buried ssDNA through transient dissociation of one or more DBDs. The second mode of ssDNA access might be provided through post-translational modifications of the DBDs or the disordered linkers(22,24). Using hydrogen-deuterium exchange mass spectrometry, we recently showed that the domains of RPA are not splayed apart but are tightly organized along with the protein-interaction domains and this configuration is altered upon phosphorylation by Aurora kinase B (22). This modification is specific to RPA functions during mitosis. To gain a better understanding of these configurational changes and how the phosphorylated RPA alters the wrapping of ssDNA, we performed crosslinking mass spectrometry (XL-MS) of RPA and phosphorylated-RPA in the absence/presence of ssDNA.

Aurora kinase B phosphorylates RPA at a single Ser-384 position in the large RPA70 subunit(22). Using genetic code expansion (33,34) we generated site-specific phospho-serine (pSer) modified RPA at Ser-384 (RPA-pSer^384^). This approach produces 100% phosphorylated RPA as seen by complete shift of the RPA-pSer^384^ band in Phos-Tag SDS-PAGE analysis (**Figure 7A**). Site-specific incorporation was also confirmed through mass spectrometry (**Supplementary Figure 5**) and by western blotting using a pSer-384 specific antibody (**Figure 7B**)(22). XL-MS of RPA or RPA-pSer^384^ were performed with a bis(sulfosuccinimidyl)suberate (BS3) crosslinker in the absence or presence of a (dT)_25_ ssDNA substrate. BS3 crosslinks primary amines (Lys residues) that are within 15 ÅÅ (42).

**Figure 7.**
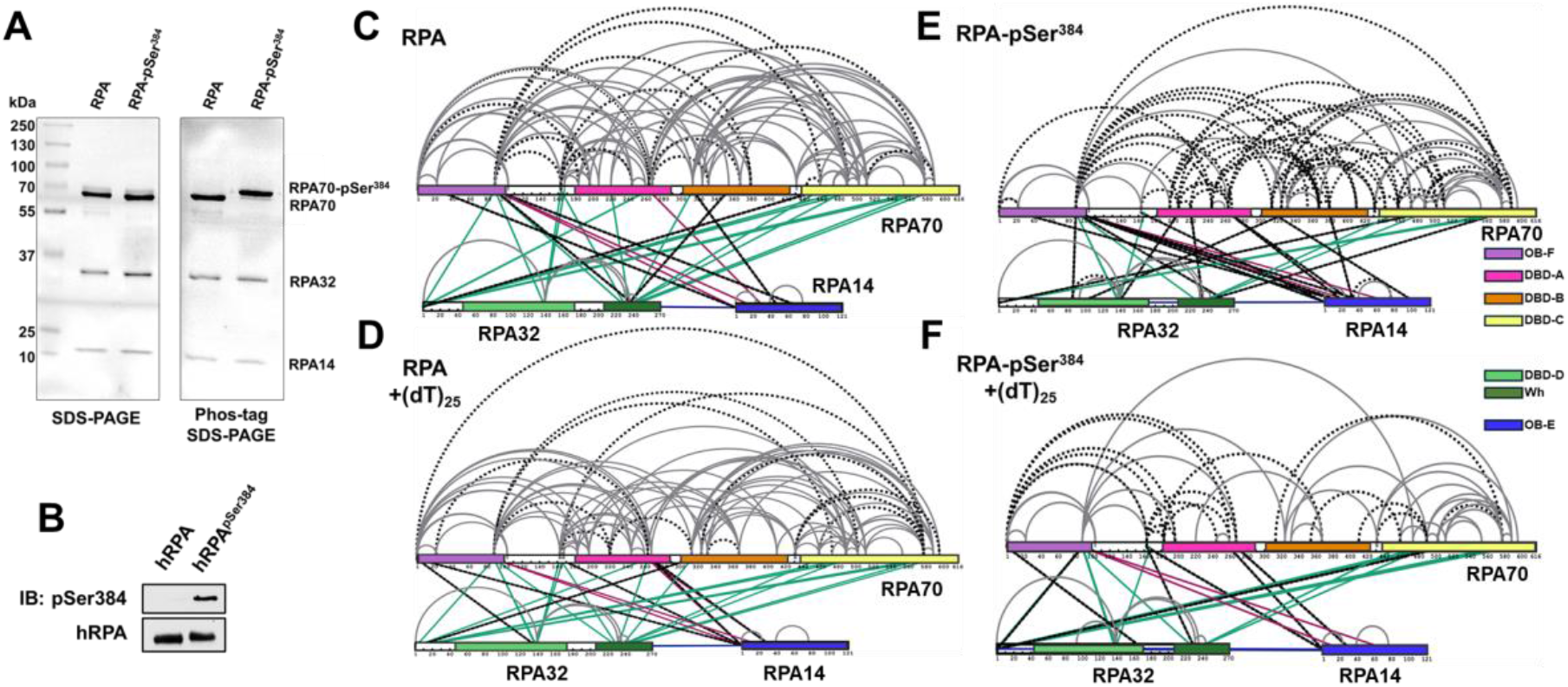
Phosphorylation at a single position in RPA70 alters access to the ssDNA with minimal alternations of end-to-end distance. **A)** SDS-PAGE and Phos-Tag SDS-PAGE analysis of RPA and RPA-pSer^384^ proteins show selective shift of the RPA70 band only in the RPA-pSer^384^ sample confirming 100% incorporation of pSer. **B)** Western blotting of RPA and RPA-pSer^384^ with an antibody specific to pSer^384^ confirms the site-specific phosphorylation. **C & D)** Cross-linking mass spectrometry (XL-MS) analysis of RPA in the absence or presence of ssDNA (dT)_25_. XLs arising from RPA32 and RPA14 are colored green and red, respectively. XLs unique to each sample are denoted by dotted lines. **F & G)** XL-MS comparison between RPA-pSer^384^ and the RPA-pSer^384^-ssDNA complex. XLs are marked in a comparative manner between tow two samples as explained in C-D.

We made several key observations: 1) Extensive crosslinks (XLs) are captured between the three subunits (RPA70, RPA32, and RPA14) and between the DNA binding domains (DBDs A, B, C, & D), the protein interactions domains (OB-F & wh), and several connecting linkers of non-phosphorylated RPA (**Figure 7C**). Of particular interest are the XLs between OB-F and F-A linker with DBD-A, DBD-B, DBD-C, DBD-D, and the wh domains. This finding suggests that all the domains are situated close together in a compacted configuration. 2) Comparison of RPA in the absence and presence of ssDNA shows crosslinks unique to each condition (denoted by the dotted lines in Figure 7), suggesting changes in the configurations/conformations or ssDNA occluded Lys residues. However, the overall contacts between the domains are still observed (**Figures 7C & D)**. These data suggest that ssDNA wraps around the compacted structural architecture of RPA without the need to unravel the domains, as would be expected if the ssDNA were to be linearly stretched. 3) Surprisingly, introduction of pSer at position 384 (DBD-B; RPA70) produces significantly new contacts within RPA (**Figure 7E & Supplemental Figure 6**). 4) Finally, the largest changes in XL patterns are observed upon ssDNA binding to RPA-pSer^384^ suggesting that a single post-translational modification can bring about large-scale configurational changes leading to an altered ssDNA-RPA-pSer^384^ complex (**Figure 7F & Supplementary Figure 7**).

To understand how ssDNA is shaped when bound to RPA-pSer^384^, we measured the end-to-end distance using DEER (**Figure 5**). We observe two distributions for RPA-pSer^384^: The first overlaps with with-type RPA and ∼70% of the phosphorylated RPA is in this population (∼7 nm). In addition, there is a 30% increase in the second population where the end-to end distance is much shorter (∼5 nm). Along with the XL-MS data, we interpret these changes as remodeling of the DBDs without large-scale changes to the wrapping pattern of the ssDNA. Thus, from a functional standpoint, phosphorylation remodels RPA such that segments of ssDNA are made accessible to RPA-interacting proteins with minimal alterations to the wrapping of DNA. For RPA-pSer^384^, these interactions are specific to mitosis.

## DISCUSSION

RPA binds ssDNA with high affinity and coats the substrate to form a regulatory protein-interaction hub to orchestrate a variety of DNA metabolic processes. RPA possesses multiple DNA binding and protein interaction domains that are connected by disordered linkers and spread across its heterotrimeric architecture. An incoming RPA-interacting protein could access the ssDNA buried under RPA by remodeling on one or more domains without the need to displace RPA. To better understand such remodeling occurs, knowledge of how ssDNA is bound by RPA is required. Canonically, models for RPA postulate that the DNA binding domains (DBDs) are assembled in a linear array to stretch the DNA leading to multiple binding modes. In contrast, structural studies show that the ssDNA is bent around the DBDs. In solution, an ensemble of states ranging from linear to the bent form can also be envisioned to exist in equilibrium.

From the perspective of the binding properties of the individual domains, recent data from our group and others support the idea of RPA behaving as two functional halves with a dynamic F-A-B half (OB-F, DBD-A, & DBD-B) and a less-dynamic Tri-C half (DBD-C, RPA32, and RPA14). Here Tri-C is modeled to provide ssDNA binding stability to RPA. Support for this model also arises from single-molecule C-trap experiments where the diffusion rate of RPA is dictated by the Tri-C half. Here, we show that ssDNA wraps around RPA and end-to-end distance measurements suggest uniform contact throughout the ssDNA-RPA complex. The data argues against linearization of ssDNA and an array-like assembly of the DBDs.

Crosslinking MS data supports a structural model for RPA where both the DBDs and protein-interaction domains (PIDs) are in close proximity. Crosslinks were also detected between the disordered OB-F:DBD-A linker and almost all DBDs and PIDs, suggesting a compacted structure for RPA (**Figure 8 & Supplementary Figure 6**). The end-to-end distance measurements, performed as a function of ssDNA length, mirror that of free ssDNA in solution with a ∼2.6 nm shift. These data suggest that ssDNA wraps around the compacted RPA core without much of a change in the intrinsic flexibility of ssDNA, likely owing to the dynamic interactions with RPA (**Figure 8**).

**Figure 8.**
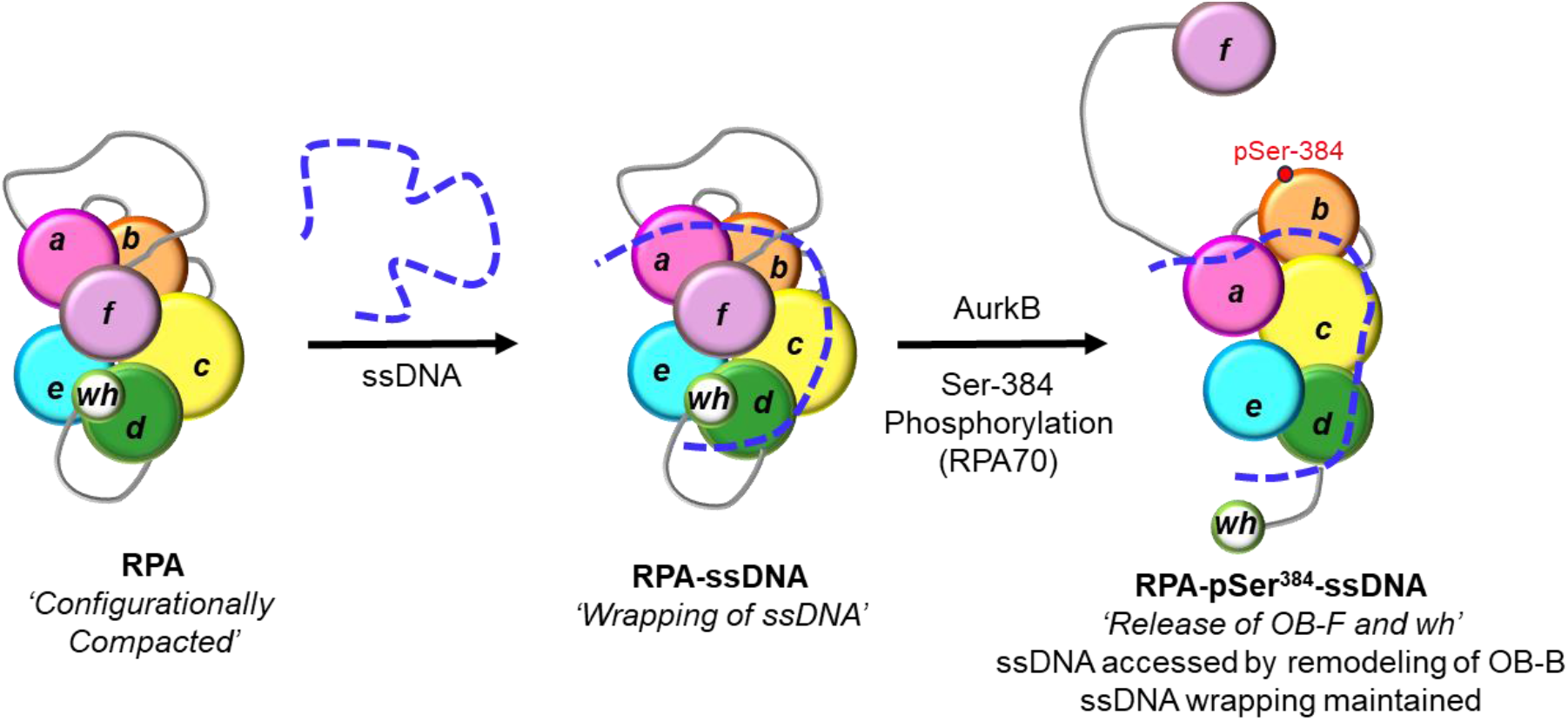
Model for ssDNA wrapping by RPA. The OB-domains of RPA are depicted along with the connecting disordered linkers. The XL-MS data suggest that the domains are compacted together and ssDNA is wrapped around this architecture. The domains are remodeled upon phosphorylation by Aurora kinase B at position Ser-384 in the RPA70 subunit. This modification releases the OB-F and wh domains and protein RPA interactions with other proteins. However, the wrapping of the ssDNA is not altered, but remodeling allows access to the ssDNA.

In such a model, access to internal regions in the ssDNA or the ends can be provided to incoming RPA-interacting proteins by rearranging one or more DBDs. Such rearrangements could be promoted by post-translational modification of one or more DBDs (**Figure 8**). As a proof of concept, we here show how phosphorylation of RPA at a single position in DBD-B (Ser-384 in RPA70) drives changes in ssDNA access. Aurora kinase B phosphorylates RPA at Ser-384 during mitosis to suppress homologous recombination and to facilitate chromosome segregation by maintaining Aurora B activity(22). Surprisingly, the end-to-end ssDNA distance does not change in RPA-pSer^384^, but the patterns of crosslinking between the domains in XL-MS are strikingly altered. Thus, the DBDs and PIDs have been repositioned or remodeled through phosphorylation without altering the overall wrapping path of ssDNA (**Figure 8**). Such changes would grant access to internal regions of the ssDNA wrapped by RPA upon phosphorylation.

In summary, measurement of end-to-end distances using smFRET and DEER spectroscopy supports a model where ssDNA is wrapped around a compacted structure of RPA with remodeling of the domains enacted through post-translational modifications. Thus, RPA can be differentially modulated to serve varying DNA metabolic needs without major changes much of a change to ssDNA organization within the complex. How these changes transpire within the context of multiple RPA molecules bound to longer ssDNA remains to be established. How ssDNA wrapping or stretching changes upon binding of multiple RPA molecules would be another interesting question to pursue, but the changes in end-to-end distances are already at the limits of detection for the smFRET and DEER methodologies described here. Thus, structural approaches such as CryoEM will need to further address this question.

## DATA AVAILABILITY

Raw data are available in the associated Supplemental Data File. Plasmids for protein overproduction as available upon request.

## SUPPLEMENTARY DATA

Supplementary Data are provided.

## AUTHOR CONTRIBUTIONS

Rahul Chadda: Formal analysis, Methodology, Validation, Writing—original draft. Vikas Kaushik: Protein Purification, pSer incorporation, Writing—review & editing. Jaigeeth Deveryshetty: Structural analysis, Writing—review & editing.

Alex Holehouse: Formal analysis, Methodology, Validation, Writing—original draft.

Iram Munir Ahmad and Snorri Th.d Sigurdsson: Design and synthesis of spin-labelled oligonucleotides, Writing—review & editing.

Reza Dastvan: EPR - Formal analysis, Methodology, Validation, Writing—review & editing.

Sofia Origanti: Conceptualization, Formal analysis, Methodology, Validation, Writing— review & editing

Edwin Antony: Conceptualization, Formal analysis, Methodology, Validation, Writing— original draft.

## ACKNOWLEDGEMENTS

Authors thank the lab members and Dr. Timothy Craggs (Exciting Instruments) for technical advice and critical reading of the manuscript. We thank Dr. Greg Sabat, University of Wisconsin-Madison, for phospho-proteomic MS analysis.

