## Supplementary Information for "Wrapping of single-stranded DNA by Replication Protein A and modulation through phosphorylation"

Supplementary Tables 1-5

Supplementary Figures 1-7

### Supplementary Table 1. Oligonucleotides used in this study

*Oligonucleotides used for the smFRET experiments*

| | |
|---|---|
| **5′ Cy5-(dT)$_{15}$-Cy3 3′** | Cy5-TTT TTT TTT TTT TTT-Cy3 |
| **5′ Cy5-(dT)$_{30}$-Cy3 3′** | Cy5-TTT TTT TTT TTT TTT TTT TTT TTT TTT TTT -Cy3 |
| **5′ Cy5-(dT)$_{45}$-Cy3 3′** | Cy5-TTT TTT TTT TTT TTT TTT TTT TTT TTT TTT TTT TTT TTT TTT TTT -Cy3 |
| **5′ Cy5-(dT)$_{60}$-Cy3 3′** | Cy5-TTT TTT TTT TTT TTT TTT TTT TTT TTT TTT TTT TTT TTT TTT TTT TTT TTT TTT TTT TTT -Cy3 |
| **5′ Cy5-(dT)$_{80}$-Cy3 3′** | Cy5-TTT TTT TTT TTT TTT TTT TTT TTT TTT TTT TTT TTT TTT TTT TTT TTT TTT TTT TTT TTT TTT TTT TTT TTT TTT TTT TT -Cy3 |
| **5′ Cy5-(dT)$_{97}$-Cy3 3′** | Cy5-TTT TTT TTT TTT TTT TTT TTT TTT TTT TTT TTT TTT TTT TTT TTT TTT TTT TTT TTT TTT TTT TTT TTT TTT TTT TTT TTT TTT TTT TTT TTT TTT T -Cy3 |

*Oligonucleotides used for the DEER experiments*

| | | |
|---|---|---|
| 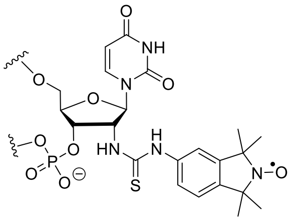 **(M)** | **(dT)$_{22}$** **DEER** | 5'-d(**M** TT TTT TTT TTT TTT TTT TTT TT **M** T)-3' |

| | | |
|---|---|---|
| 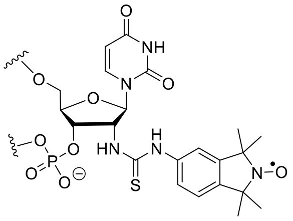 **(M)** | **(dT)$_{50}$** **DEER** | 5'-d (TTT **M**TT TTT TTT TTT TTT TTT TTT TTT TTT TTT TTT TTT TTT TTT TTT TTT **M**T) 3' |

**Supplementary Table 2. Composition of media and reagents used for Phosphoserine incorporation media.**

**A) ZY-non inducing media (ZY-NIM)**

| **Components** | **For 50 mL** |
|---|---|
| ZY media | 47.25 mL |
| 25x M-salts | 0.1 mL |
| 1 M $MgSO_4$ | 2 mL |
| 40 % (w/v) α-D-glucose | 0.625 mL |
| Trace metals solution (5000x) | 0.01 mL |

**B) ZY-auto inducing media (ZY-AIM)**

| **Components** | **For 1 L** |
|---|---|
| ZY media | 940 mL |
| 1 M $MgSO_4$ | 2 mL |
| 25x M-salts | 40 mL |
| 50x 5052 solution | 20 mL |
| Trace metals solution (5000x) | 0.2 mL |

**C) Composition of 25X M-salts**

| **Components** | **For 1 L** |
|---|---|
| Sodium phosphate dibasic | 88.73 g |
| Potassium phosphate dibasic | 85.05 g |
| Ammonium chloride | 66.86 g |
| Sodium sulfate anhydrous | 17.75 g |

**D) 50x 5052 solution**

| **Components** | **For 1 L** |
|---|---|
| α-D-glucose | 2.5 g |
| Lactose | 50 g |
| Glycerol (v/v) | 125 mL |

**Supplementary Table 3. Comparison of FRET efficiencies measured using the EI-FLEX and the Picoquant MT200**

| | FRET efficiency | | Nbursts | |
|---|---|---|---|---|
| | **EI-FLEX** | **MT200** | **EI-FLEX** | **MT200** |
| $(dT)_{15}$ | 0.83 (±0.064) | 0.89 (±0.11) | 765 | 10260 |
| $(dT)_{30}$ | 0.37 (±0.086) | 0.426 (±0.097) | 268 | 9056 |
| $(dT)_{45}$ | 0.15 (±0.060) | 0.17 (±0.088) | 366 | 11788 |
| $(dT)_{60}$ | 0.074 (±0.049) | 0.066 (±0.078) | 511 | 18453 |
| $(dT)_{80}$ | 0.021 (±0.05) | 0.022 (±0.075) | 426 | 40425 |
| $(dT)_{97}$ | 0.027 (±0.37) | 0.032 (±0.088) | 1257 | 636 |

**Supplementary Table 4. FRET efficiencies measurements (smFRET)**

| **$(dT)_{25}$ and hRPA** | **FRET efficiency** | | |
|---|---|---|---|
| | **Free** | **Bound** | **N bursts** |
| $(dT)_{25}$ | 0.49 (±0.093) | - | 1072 |
| $(dT)_{25}$ + 0.02 nM hRPA | 0.49 (±0.095) | 0.07 (±0.032) | 1086 |
| $(dT)_{25}$ + 0.1 nM hRPA | 0.49 (±0.11) | 0.075 (±0.047) | 852 |
| $(dT)_{25}$ + 1 nM hRPA | - | 0.07 (±0.2) | 912 |

| **$(dT)_{25}$ and yRPA** | **FRET efficiency** | | |
|---|---|---|---|
| | **Free** | **Bound** | **N bursts** |
| $(dT)_{25}$ | 0.49 (±0.093) | - | 1072 |
| $(dT)_{25}$ + 0.02 nM ScRPA | 0.48 (±0.098) | 0.09 (±0.031) | 773 |
| $(dT)_{25}$ + 0.1 nM ScRPA | 0.47 (±0.11) | 0.089 (±0.085) | 783 |
| $(dT)_{25}$ + 1 nM ScRPA | - | 0.091 (±0.12) | 1172 |

**Supplementary Table 5. FRET efficiencies measurements (smFRET)**

| **$(dT)_{30}$ and hRPA** | **FRET efficiency** | | |
|---|---|---|---|
| | **Free** | **Bound** | **N bursts** |
| $(dT)_{30}$ | 0.37 (±0.098) | - | 585 |
| $(dT)_{30}$ + 0.03 nM hRPA | 0.38 (±0.095) | 0.04 (±0.055) | 654 |
| $(dT)_{30}$ + 0.1 nM hRPA | 0.37 (±0.092) | 0.025 (±0.068) | 568 |
| $(dT)_{30}$ + 1 nM hRPA | - | 0.055 (±0.047) | 569 |

| **$(dT)_{45}$ and hRPA** | **FRET efficiency** | | |
|---|---|---|---|
| | **Free** | **Bound** | **N bursts** |
| $(dT)_{45}$ | 0.17 (±0.075) | - | 1108 |
| $(dT)_{45}$ + 0.03 nM hRPA | 0.17 (±0.081) | 0.02 (±0.044) | 773 |
| $(dT)_{45}$ + 0.1 nM hRPA | 0.17 (±0.059) | 0.021 (±0.033) | 779 |
| $(dT)_{45}$ + 1 nM hRPA | - | 0.02 (±0.035) | 677 |

**Supplementary Table 6. DEER measurements**

| | **<*R*> [nm]** | **$\sigma$ [nm]** | ***f* (%)** |
|---|---|---|---|
| **$(dT)_{22}$** | 3.1±1.0 | 0.7±0.6 | 36 |
| | 5.1±2.0 | 1.1±3.1 | 64 |
| **$(dT)_{22}$+hRPA** | 4.6±0.4 | 1.0±0.7 | 5 |
| | 6.8±0.8 | 0.4±1.2 | 95 |
| **$(dT)_{22}$+hRPA$^{pSer384}$** | 4.6±0.4 | 1.0±0.7 | 30 |
| | 6.8±0.8 | 0.4±1.2 | 70 |
| **$(dT)_{22}$+yRPA** | 6.9±0.3 | 1.0±0.3 | 100 |

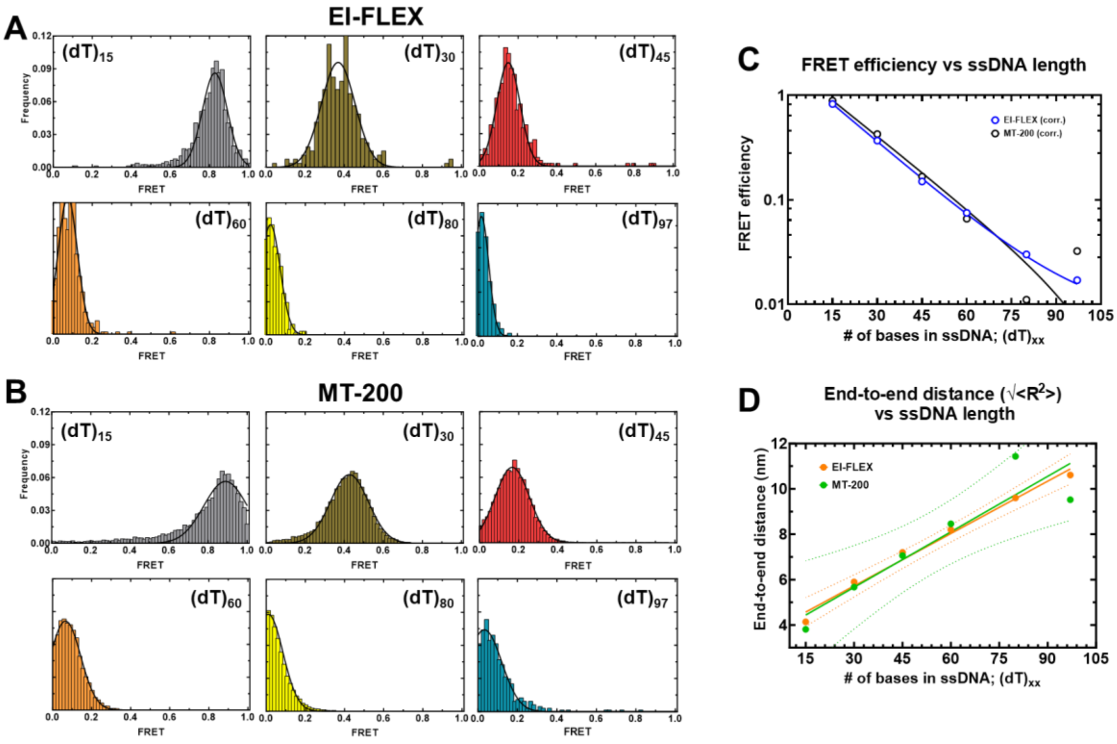


**Supplementary Figure 1. Comparison of smFRET data collected using the EI-FLEX and MT-200 microscopes.** smFRET data collected for various lengths of ssDNA using the **A)** EI-FLEX and **B)** Picoquant MT-200 microscopes. **C)** Plot of the FRET efficiencies versus ssDNA shows excellent agreement between measurements made using both instruments. **D)** Similarly, end-to-end distance measurements calculated from the smFRET data show excellent agreement for both instruments.

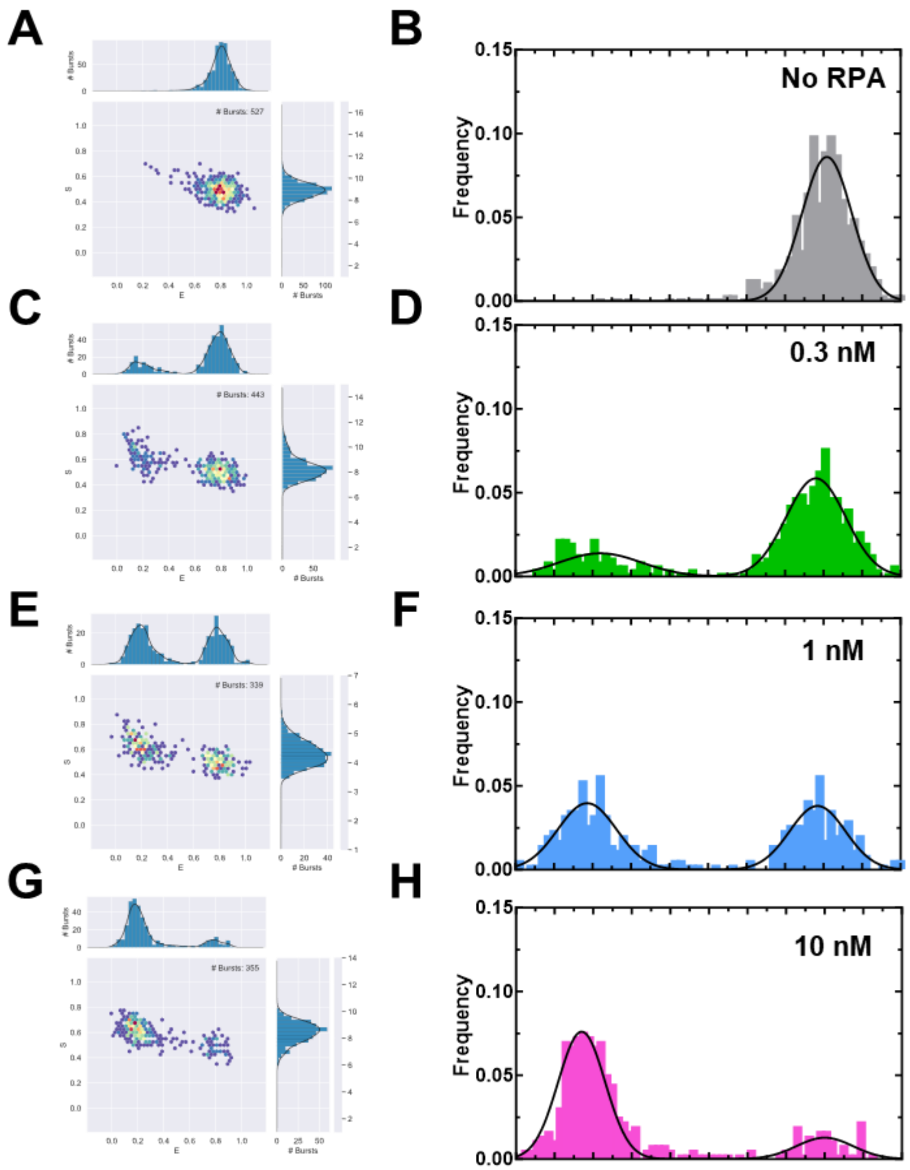


**Supplementary Figure 2. smFRET data for RPA-(dT)15 complexes.** smFRET data collected for the (dT)15 substrate in the absence of RPA **(A & B)** and increasing concentrations of RPA **(C-H)**.

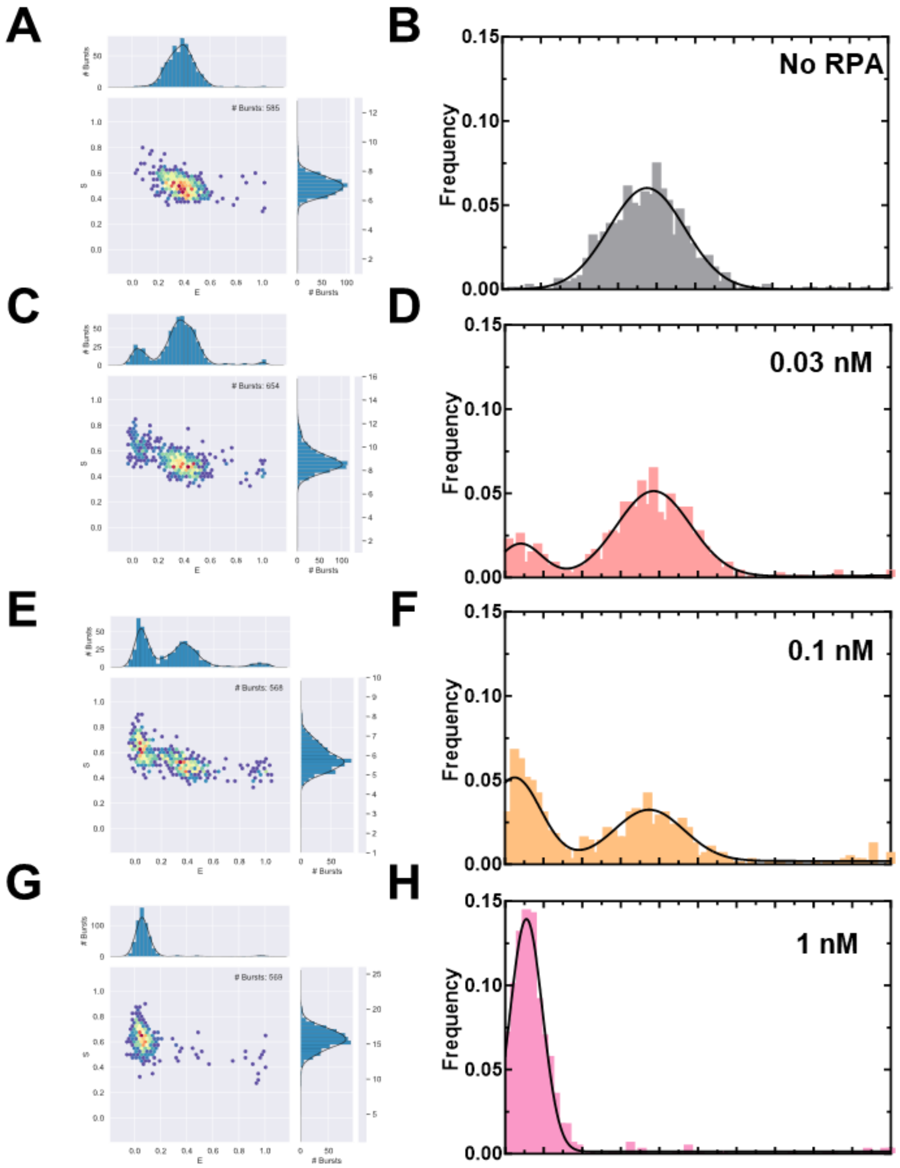


**Supplementary Figure 3. smFRET and mass photometry data for RPA-(dT)$_{30}$ complexes.** smFRET data collected for the (dT)$_{30}$ substrate in the absence of RPA **(A & B)** and increasing concentrations of RPA **(C-H)**.

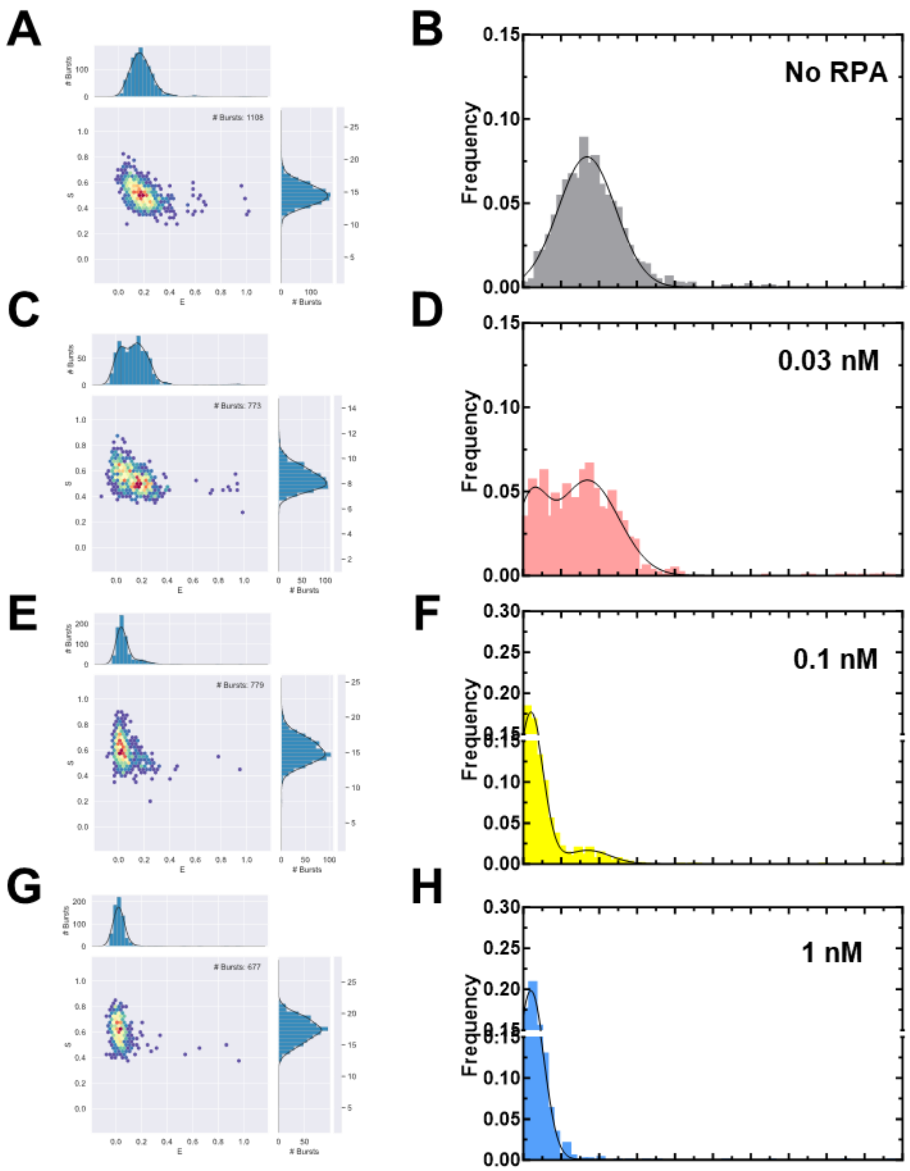


**Supplementary Figure 4. smFRET and mass photometry data for RPA-(dT)$_{45}$ complexes.** smFRET data collected for the (dT)$_{45}$ substrate in the absence of RPA **(A & B)** and increasing concentrations of RPA **(C-H)**.

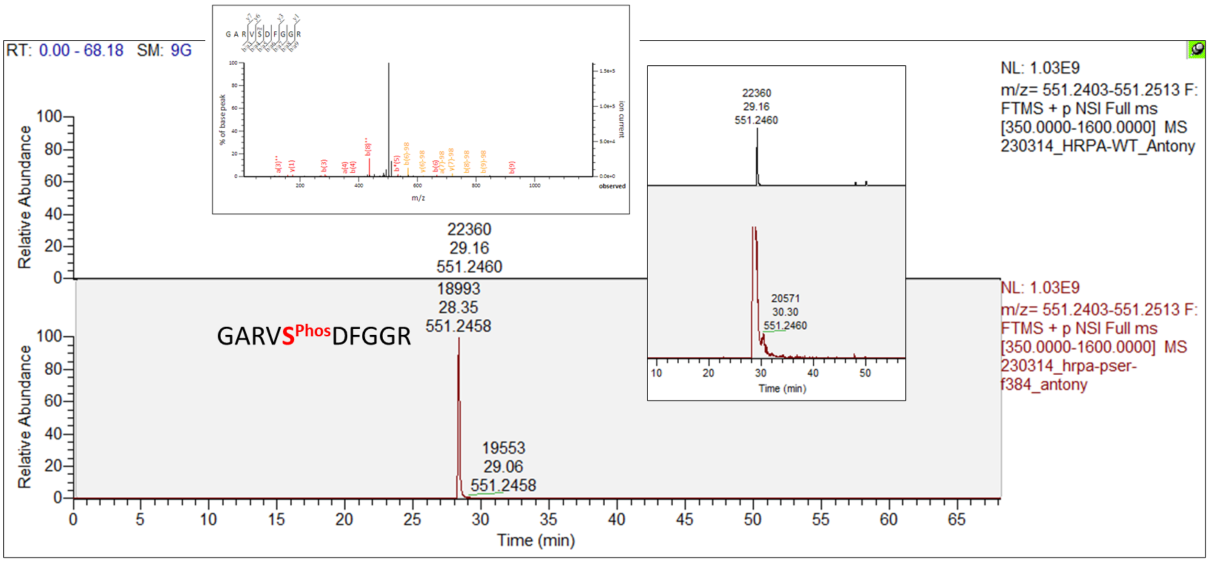


**Supplementary Figure 5. Mass spectrometry analysis of pSer incorporation in hRPA.** Mass spectrometry analysis of RPA-pSer[384] shows site-specific phosphoserine incorporation.

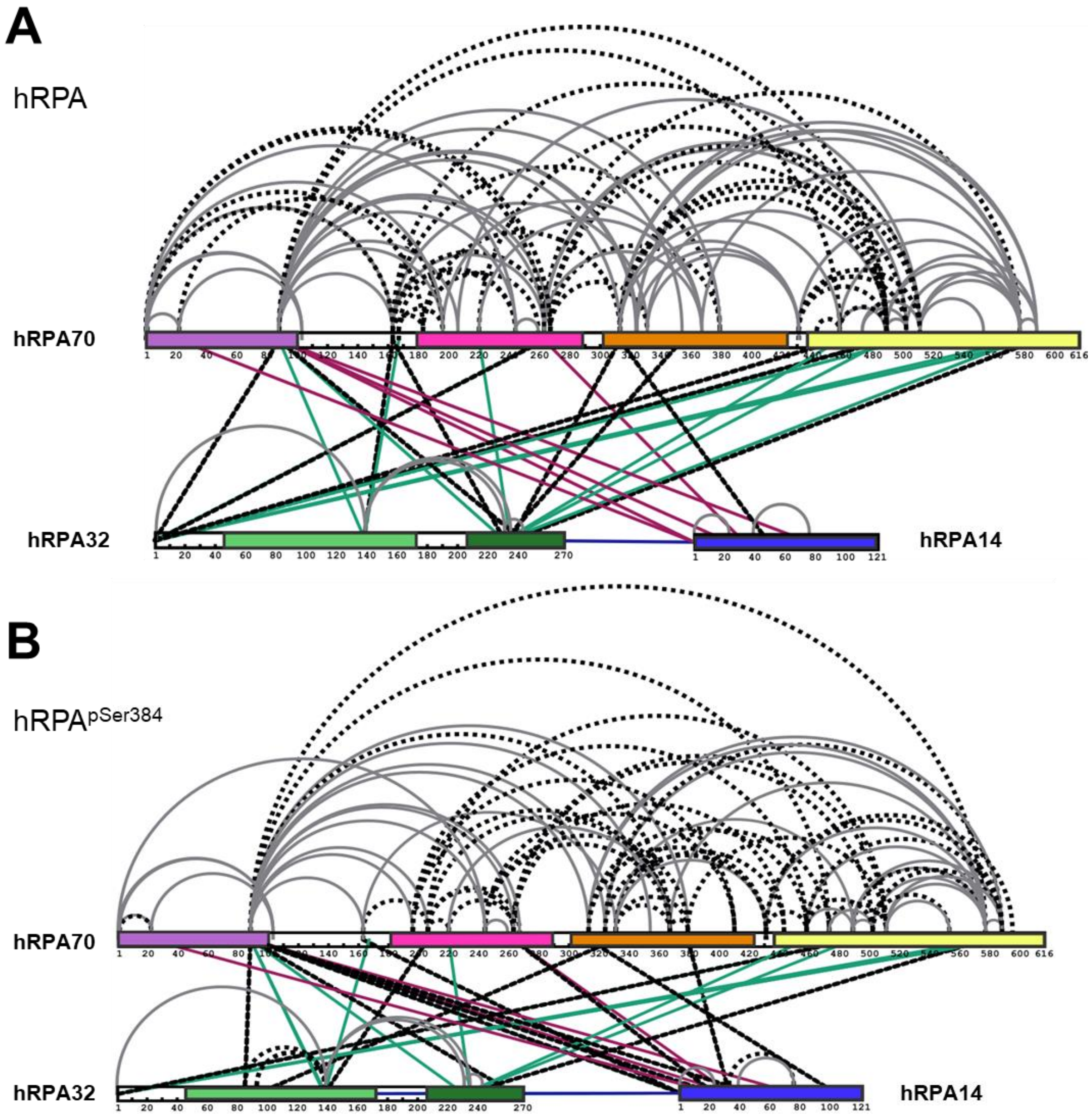


**Supplementary Figure 6. Unique XLs in the RPA introduced by phosphorylation at Ser-384 in RPA70.** Crosslinking mass spectrometry (XL-MS) analysis of **A)** RPA and **B)** RPA-pSer$^{384}$ are shown. The two datasets are compared relative to each other and the crosslinks unique to each sample are shown in dotted black lines. Crosslinks in grey are common to both datasets and depict intra-subunit crosslinks within RPA70-, RPA32 and RPA14. Inter-subunit crosslinks between RPA70, RPA32, and RPA14 are shown in green and red, respectively.

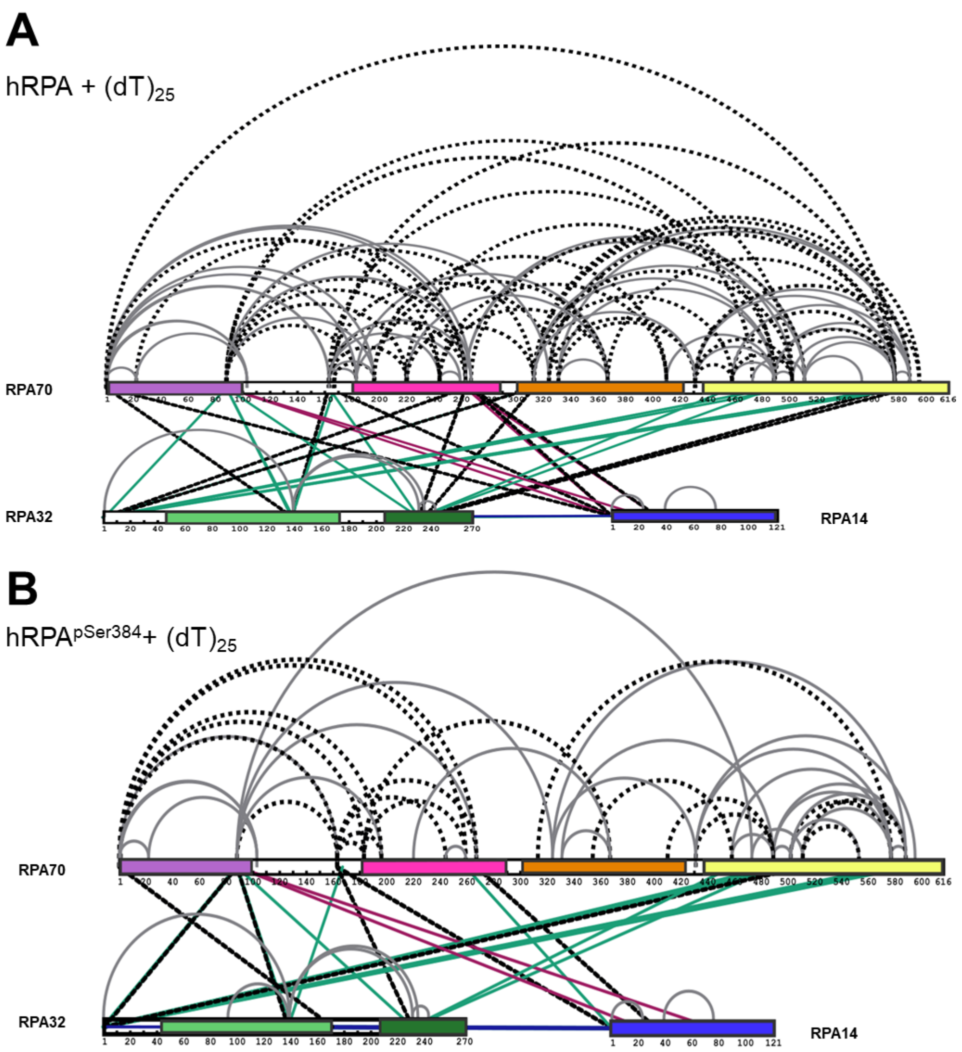


**Supplementary Figure 7. Unique XLs in the RPA-ssDNA complex introduced by phosphorylation at Ser-384 in RPA70.** Crosslinking mass spectrometry (XL-MS) analysis of **A)** RPA-ssDNA and **B)** RPA-pSer$^{384}$-ssDNA complexes are shown. The two datasets are compared relative to each other and the crosslinks unique to each sample are shown in dotted black lines. Crosslinks in grey are common to both datasets and depict intra-subunit crosslinks within RPA70-, RPA32 and RPA14. Inter-subunit crosslinks between RPA70, RPA32, and RPA14 are shown in green and red, respectively.
